# Ionizable Lipids Promote Curvature Remodeling and Altered Fluctuation Dynamics in Endosomal Membranes

**DOI:** 10.64898/2026.08.17.745287

**Authors:** Teshani Kumarage, Yiding Li, Bengu Sueda Sengul, Mayesha B. Mustafa, Jinchao Lou, Michael D. Best, Charles M. Schroeder, Cecilia Leal

**Affiliations:** Department of Materials Science and Engineering, The Grainger College of Engineering, University of Illinois Urbana-Champaign, Urbana, IL 61801, USA; Department of Chemistry, University of Tennessee, 1420 Circle Drive, Knoxville, TN 37996, USA; Department of Chemistry & Biochemistry, University of Mississippi, MS 38677, USA

## Abstract

Inefficient endosomal escape is a crucial barrier to intracellular delivery of nucleic acid therapeutics using lipid nanoparticles (LNPs). The use of ionizable lipids (ILs) has significantly improved cargo delivery efficiency, yet the physical basis of their interaction with endosomal membranes and their role in endosomal escape remain unclear. It has been suggested that, as ILs become cationic during endosomal acidification, electrostatic affinity promotes fusion of the LNPs with the endosome. In this paper, we propose an additional mechanism in which ILs are redistributed from LNPs to host membranes, modulating the elastic properties and curvature of the membrane, lowering the energetic threshold for endosome disruption. To test this, we quantified the spontaneous curvature of clinically relevant ILs and ATP-binding lipids and measured the membrane mechanics of giant unilamellar vesicles (GUVs) with an endosome-relevant composition at endosome-relevant pH. Small-angle X-ray scattering (SAXS) measurements reveal that the incorporation of ILs and ATP lipids into endosome-mimetic membranes shifts the spontaneous curvature towards more negative values. Micropipette aspiration experiments indicated a decrease in the apparent area compressibility modulus of membranes doped with ILs and ATP lipids. In addition, membranes showed enhanced fluctuation amplitudes and altered relaxation behavior, consistent with membrane perturbations associated with lipid insertion and pH- or ATP-driven destabilization. Under conditions promoting the partitioning of ILs or ATP-binding lipids, we further observed reduced bending rigidity and increased heterogeneity in membrane tension. Together, these results support a model in which ILs (as well as newly developed ATP-binding lipids) partition into endosomal membranes, softening the membrane and generating local curvature frustration that facilitates endosomal disruption during the natural acidification process. By quantitatively linking lipid composition with changes in membrane elasticity and fluctuation dynamics, this work provides a biophysical framework for understanding how lipid redistribution may contribute to endosomal escape and improve delivery efficiency.

**SIGNIFICANCE:** Efficient endosomal escape remains a major bottleneck to the success of LNP-mediated nucleic acid delivery. Despite the central role of ionizable lipids (ILs), how they interact with the endosomal membrane and promote destabilization remains unclear. Here, we provide biophysical evidence that IL incorporation remodels the mechanical properties of endosome-mimetic membranes. By quantifying lipid spontaneous curvature together with membrane tension and fluctuation dynamics, we show that IL incorporation promotes negative curvature, softens membranes, and enhances fluctuations. These changes may lower the energetic barrier for membrane deformation and disruption required for endosomal escape. Our findings offer a quantitative link between lipid properties and membrane mechanics, providing insight into how ILs promote endosomal escape and informing design principles for improving LNP-based therapeutics.

## INTRODUCTION

Messenger RNA (mRNA) therapeutics are a promising class of medicine that enables transient, programmable protein expression without altering the genome. Since the discovery of its role as the intermediary between DNA and proteins in the mid-20th century (1), mRNA has evolved into a therapeutic platform through advances in transcription *in vitro* (2) and *in vivo* (3), yet with significant challenges such as instability and immunogenicity. In recent decades, innovations in RNA chemistry (4, 5) and lipid nanoparticle (LNP) delivery systems (6) have overcome some of these barriers, culminating in the clinical success of mRNA vaccines and opening the door to RNA treatments more generally for a wide range of diseases (6–11). These successes underscore the potential of LNPs as versatile carriers for nucleic acid (NA)-based therapeutics, capable of protecting RNA from degradation, facilitating cellular uptake, and enabling cytosolic delivery. Their efficacy, however, hinges on a critical step: the escape of therapeutic payloads from endosomes into the cytosol (12–15). Although LNPs are efficiently internalized via endocytosis, their cargo is exposed to the cell’s natural endosomal maturation pathway, in which early endosomes progressively acidify and transition into late endosomes and lysosomes containing nucleases and other hydrolytic enzymes, ultimately leading to rapid loss of RNA unless escape to the cytosol occurs (16–18).

Despite the advances in LNP design, the mechanisms governing endosomal escape remain poorly understood. LNPs need to rapidly promote functional instabilities of the endosomal membranes through which their cargo can be transported into the cytosol. These membrane instabilities typically involve fusion and/or rupture. Several recent studies demonstrate that endosomal-escape efficiency can be modulated by changing LNP lipid composition, particularly through the introduction of ionizable lipids (ILs), as well as by altering LNP nanostructure (6, 7, 14, 19–31). Our research has demonstrated that the use of *Gaussian curvature lipids* is beneficial to the delivery of mRNA (14, 15) and siRNA (26, 27, 29) because they promote non-lamellar LNP nanostructures that drive fusion processes with endosomal membranes and fusion-pore formation to release cargo. Current models propose that endosomal acidification protonates ILs, increasing their electrostatic interactions with anionic lipids in the endosomal membrane and promoting membrane fusion and cargo release into the cytosol (6). In contrast, recent studies have demonstrated that state-of-the-art LNPs containing 50 mol % of ILs mediate endosomal escape via catastrophic rupture of the endosomal membrane, leading to galectin recruitment and a concomitant inflammatory process (32). Therefore, ILs represented an important advance in the development of LNPs for mRNA vaccines, but their use as a more generic delivery platform raises concerns of immunogenicity (33, 34). Although these models emphasize interactions between the LNP and endosomal membrane, we propose an additional mechanism in which LNP-derived lipids redistribute into the endosomal membrane and alter its intrinsic biophysical properties, including curvature and elasticity, thereby increasing its susceptibility to disruption.

Figure 1 displays the chemical structures of the most commonly used ILs in LNPs. At neutral pH, ILs are uncharged, but under acidic conditions, such as those in the endosomal lumen, their headgroups become protonated. This pH-responsive behavior promotes favorable electrostatic interactions with negatively charged endosome components and is thought to contribute to membrane destabilization and endosomal escape (6, 35). However, beyond increasing electrostatic interactions with anionic membrane components, protonation can alter lipid packing and molecular shape or conformation, particularly given the bulky hydrophobic tails of many ILs. How these molecular changes affect the curvature and mechanical properties of endosomal membranes remains poorly understood. In recent years, there is growing evidence that the biophysical properties of endosomal membranes themselves, such as membrane tension, stiffness, and curvature, also regulate endosomal escape (36, 37). However, the precise biophysical mechanisms through which different LNP lipids, including ionizable, cationic, and nonlamellar lipids, interact with endosomes and what type of instabilities they drive are incompletely understood.

**Figure 1:**
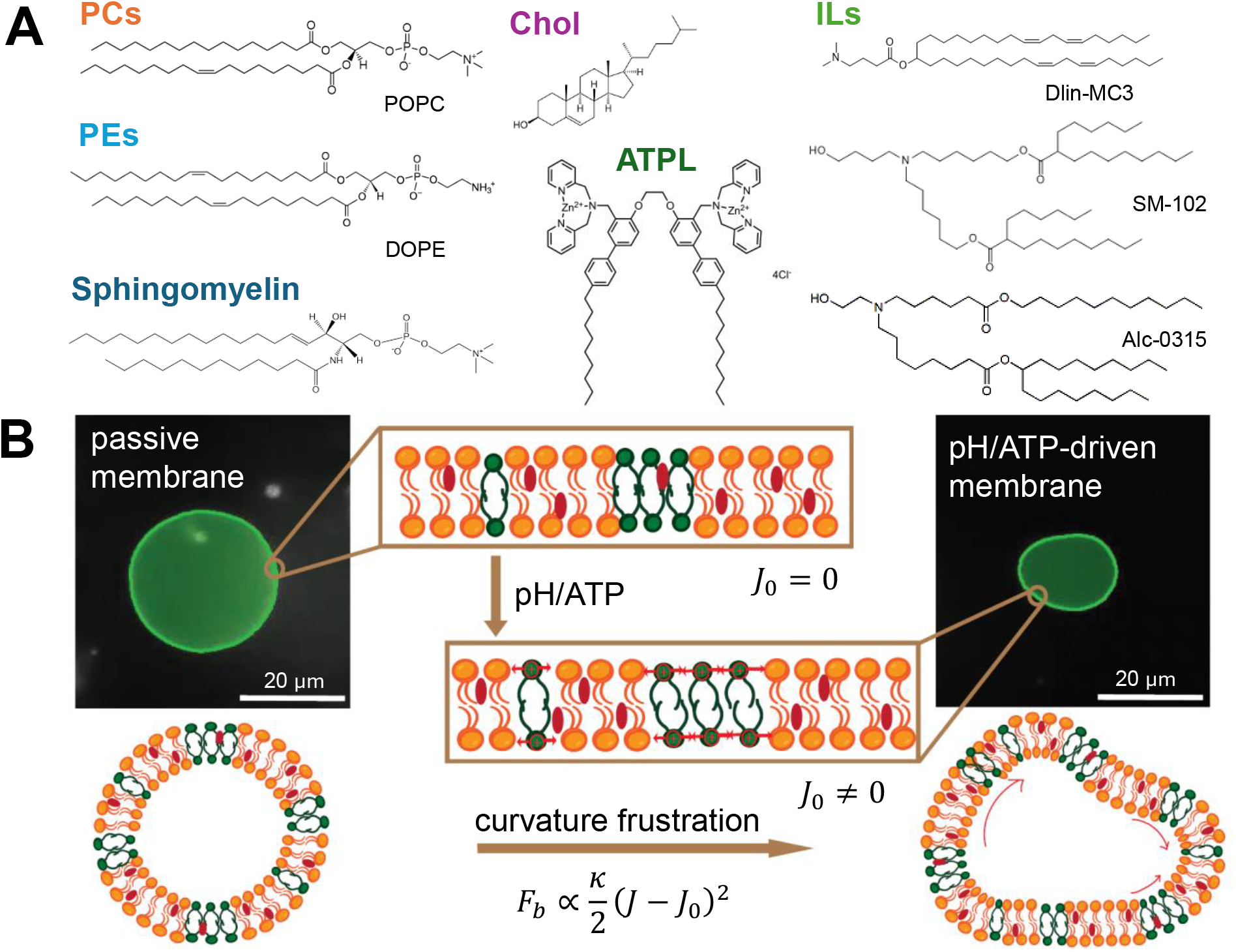
(A) Chemical structures of the lipids used in this work. Representative membrane lipids include the phosphatidylcholine (PC) lipid POPC, the phosphatidylethanolamine (PE) lipid DOPE, sphingomyelin (SM), and cholesterol (Chol). Ionizable lipids (ILs) commonly employed in LNP formulations are represented by Dlin-MC3-DMA (Dlin-MC3), SM-102, and Alc-0315. The ATPL represents a recently developed ATP-responsive lipid containing zinc-coordinating headgroups that interact with phosphate-containing species, including ATP. (B) Schematic illustrating how ILs/ATPL modulate membrane mechanics. Under steady-state conditions (left), membranes containing neutral ILs exhibit near-zero curvature (*J*_0_ ≈ 0) resulting in a stable, quasi-spherical vesicle. Upon activation by acidic pH or ATP binding (right), protonation or ligand binding alters lipid packing and increases headgroup area, shifting the preferred curvature toward negative values (*J*_0_ ≠ 0). Because the global vesicle geometry constrains large-scale deformation, this curvature mismatch generates curvature frustration, increasing the bending energy. The membrane accommodates this stress through enhanced local deformations and dynamic fluctuations rather than global shape changes, as illustrated by the distorted vesicle contour.

In this study, we use small-angle X-ray scattering (SAXS) methods to quantitatively estimate the spontaneous curvature *J*_0_ of different ILs used in LNP design: Dlin-MC3, SM-102, and Alc-0315 at neutral and acidic pH, and further evaluate how their incorporation changes the elastic properties of giant unilamellar vesicles (GUVs) mimicking early-endosomal membrane (EEM) composition via vesicle fluctuation analysis. In addition, we compare the results obtained for ILs to a new class of ATP-responsive lipids (ATPL) to recreate the conditions associated with biological endosomal acidification, a process powered by ATP consumption. ATPL contain zinc-coordinating headgroups (38) (Figure 1A) that interact with phosphate-containing species such as ATP, inducing a change in molecular packing that has the potential to destabilize endosomal membranes. We found that the spontaneous curvature of ILs and ATPL is very responsive to pH changes and ATP addition, respectively. As a result, the incorporation of these responsive lipids into EEMs leads to enhanced fluctuations and softer membranes, thereby increasing susceptibility to membrane destabilization and potentially facilitating endosomal escape (Figure 1B). By highlighting how LNP-derived lipids have the ability to directly remodel the biophysical properties of early endosomal membranes, our findings broaden the mechanistic understanding of endosomal escape and suggest new design principles for next-generation LNP delivery systems.

## MATERIALS AND METHODS

### Materials

POPC (1-Palmitoyl-2-oleoyl-sn-glycero-3-phosphocholine), DOPE (1,2-dioleoyl-sn-glycero-3-phosphoethanolamine), cholesterol, brain sphingomyelin (SM), and Rhodamine PE (1,2-dioleoyl-sn-glycero-3-phosphoethanolamine-N-(lissamine rhodamine B sulfonyl)) were purchased from Avanti Polar Lipids (AL, USA) and used without further purification. The ILs DLin-MC3, SM-102, and Alc-0315 were purchased from Cayman Chemicals (MI, USA) and used without further purification. ATPL was synthesized at the University of Tennessee, Knoxville (38) (Figure 1A).

### Sample preparation for fluctuation analysis and micropipette aspiration

Giant unilamellar vesicles (GUVs) were prepared using the electroformation protocol with indium-tin-oxide (ITO)-covered slides (39). GUVs prepared with early endosomal (EEM) composition consisted of 40 mol% POPC, 20 mol% DOPE, 6 mol% SM, and 34 mol% cholesterol (40, 41) with 0.3 mol% of Rhodamine PE. First, lipids were dissolved in chloroform and dried under a gentle flow of *N*_2_ to form a uniform thin film, then dried under vacuum overnight. After all residual chloroform was removed, ILs (dissolved in EtOH) were added, mixed by vortexing, and dried overnight. The thin film was then redissolved in chloroform again and well mixed by vortexing to ensure a homogeneous mixture. The prepared solution (10 *µ*L) was transferred onto thoroughly cleaned and dried ITO plates and uniformly spread with 2.5 *µ*L methanol using a 22G syringe needle. The ITO plates were then subjected to overnight drying under vacuum. Finally, the films were hydrated in sucrose solution (100 mM, pH 7.4) and the electrodes were connected to an AC power supply (10 Hz, 20 *V*_pp_) at 55 °C for at least 3 hrs. In this work, we prepared GUVs with EEM membrane composition along with 10 mol% ILs/ATPL and 0.3 mol% Rhodamine PE. The ILs used in this study, Dlin-MC3, SM-102, and Alc-0315, have 2, 3, and 4 hydrocarbon chains, respectively (Figure 1). For the control experiments, the GUV samples were used as-is, and to mimic acidic conditions, sodium acetate (NaOAc) buffer (pH 5.5) was added to the GUV sample in a 4:1 ratio. In the case of ATPL, a 1 mM ATP solution was introduced in the same ratio.

### Sample preparation for SAXS

All samples used for SAXS experiments were prepared as bulk lipid mixtures. The required amounts of lipids were dissolved in chloroform and dried under a gentle stream of *N*_2_ to form a thin lipid film, then dried under vacuum overnight. IL, dissolved in ethanol, was introduced to the vial and dried under the same conditions to remove residual solvent. The thin film was then redissolved in chloroform and vortex-mixed thoroughly to ensure a homogeneous mixture was produced. The lipid mixture was subsequently transferred into quartz capillaries (Hilgenberg, Germany) and allowed to air dry for 2-3 days, followed by additional vacuum drying to ensure complete removal of chloroform. After solvent evaporation, the samples were hydrated with either Milli-Q water or NaOAc buffer (pH 5.5). The capillaries were centrifuged at 3000 rpm for 3 minutes to facilitate sample settling and then allowed to hydrate for 1-2 days. Prior to measurements, capillaries were flame sealed and further secured with epoxy to prevent sample dehydration during storage and data collection.

### Spontaneous curvature from SAXS

Preliminary SAXS experiments were performed using the in-house SAXS instrument custom-built in collaboration with Forvis Technologies (Santa Barbara, CA, USA). The system was equipped with a Xenocs GeniXD ultralow-divergence Cu *K*_*α*_ X-ray source (8.0 keV) with a beam divergence of 1.3 mrad. The sample-to-detector distance and scattering vector *q* calibration were determined using a silver behenate (AhBeH) standard. 2D diffraction patterns were acquired using a Pilatus 300K 20 Hz hybrid pixel detector (Dectris). The resulting 2D diffraction images were reduced to a 1D scattering profile using Fit2D software developed by the European Synchrotron Radiation Facility (ESRF). Synchrotron SAXS experiments were performed at beamline 12-ID-B of the Advanced Photon Source (APS) at Argonne National Laboratory using a 13.3 keV X-ray beam. Scattering patterns were collected using a Pilatus 2M detector (Dectris) with a pixel size of 0.172 mm. The acquired 2D SAXS data were radially integrated on-site using MATLAB software developed at beamline 12-ID-B. The sample-to-detector distance was calibrated using an AgBEH standard. The spontaneous curvature (*J*_0_) was estimated from the temperature dependence of the inverse hexagonal (*H*_II_) phase lattice parameters obtained by SAXS, following the approach of Gillams *et al*. (42). In brief, at each temperature, the lattice parameter *a* was determined from the SAXS profile; *a* represents the center-to-center spacing between adjacent water cylinders. For the *H*_II_ phase, *a* is related to the water core radius *R*_*w*_ and lipid monolayer thickness *l* through the relation:

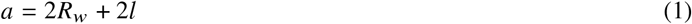

where *l* was taken to be 1.48 nm for DOPE. The monolayer curvature is then approximated from the radius at the pivotal plane,

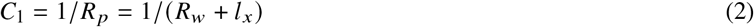

where *l*_*x*_ is the distance between the pivotal plane and the water interface, assumed to be weakly temperature dependent. To determine the spontaneous curvature, the lattice parameters were measured as a function of temperature, allowing principal curvature values *C*_1_ to be calculated at different temperatures.

The elastic properties of membranes are strongly influenced by their spontaneous curvature *J*_0_, as described by the Helfrich elastic formalism. In this framework, the membrane bending energy depends on the deviation of the total curvature *J* = *C*_1_ + *C*_2_ from the spontaneous curvature, as well as on the Gaussian curvature *K* = *C*_1_ *C*_2_. The energetic contributions associated with these deformations are weighted by the bending modulus *κ* and the Gaussian modulus 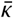, respectively (43, 44) via the relation:

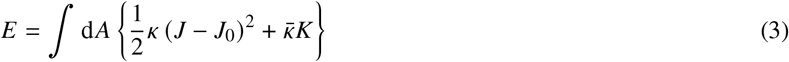

Using this theory, the difference in the curvature elastic energy between two temperatures can be rearranged into a linear form:

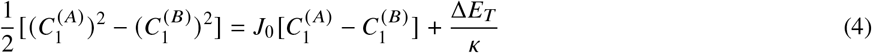

Thus, plotting 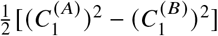 versus 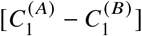 yields a straight line, whose slope gives the spontaneous curvature of the mixture 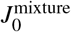. It is important to note that this approach assumes similar bending rigidities of mixture components and negligible temperature dependence of structural parameters such as the pivotal plane position. For well-mixed lipid systems, the spontaneous curvature can be approximated as a mole-fraction-weighted (Φ) sum of the spontaneous curvatures of the individual lipid species (45, 46):

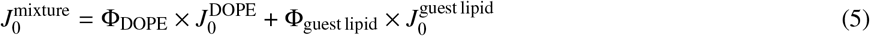

This condition was achieved through mixing of the lipids in chloroform solution before solvent removal, assuming the absence of phase separation.

### Vesicle segmentation and contour tracking

To determine membrane contours from time-lapse fluorescence microscopy recordings, we employed an automated segmentation pipeline based on the Segment Anything Model 2 (SAM2) (47), a deep learning model developed for image and video segmentation. In the first frame of each video, the vesicle was identified by manually placing one or more marker points within the membrane. SAM2 then automatically propagated this identification through all subsequent frames, generating a binary segmentation mask *M* (*x, y, t*) that distinguishes the vesicle from the background in each frame. From each mask, the vesicle boundary was extracted as a closed contour using standard image processing routines (OpenCV) (48), preserving the full pixel-resolution detail of the membrane edge. When multiple regions were detected, the largest contour by area was retained. The resulting contour coordinates (*x*_*i*_, *y*_*i*_) were exported for each frame and used as input for fluctuation analysis. This approach is robust to intensity variations and photobleaching and maintains consistent tracking through minor defocusing events during imaging, allowing for minimal manual intervention beyond the initial annotation.

### Vesicle fluctuation analysis

To characterize the mechanical properties of GUVs, we performed fluctuation analysis based on the refined contour detection methodology developed by Pécréaux *et al*. (49). Time-lapse imaging of GUV equatorial cross-sections was used, and membrane contours were determined from each frame as discrete coordinate sets (*x*_*i*_, *y*_*i*_). Contour quality was assessed by the arc length, computed via periodic cubic spline fitting with smoothing parameter *s* = *Nϵ*^2^, where *N* is the number of contour points and *ϵ* ≈ 0.5 pixels represents the subpixel localization uncertainty. This smoothing suppresses pixel-scale artifacts that otherwise inflate the measured contour length. Contours deviating by more than 10% from the mean length were discarded (49). The center of mass of each contour was computed using arc-length-weighted averaging, and centered coordinates were transformed to polar coordinates (*r*_*i*_, *θ*_*i*_). The polar contour was then decomposed into discrete Fourier modes:

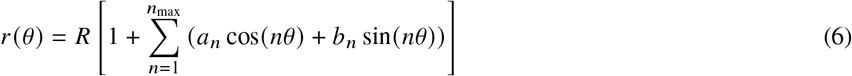

where *R* is the mean radius and the Fourier coefficients *a*_*n*_ and *b*_*n*_ were calculated using trapezoidal integration to accommodate non-uniform angular spacing. The fluctuation spectrum was obtained from the variance of the complex Fourier coefficient *c*_*n*_ = *a*_*n*_ − *ib*_*n*_:

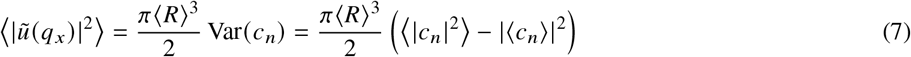

where *q*_*x*_ = *n*/⟨*R*⟩ is the wave vector corresponding to mode number *n*, and the variance is computed over all frames in the time series. The experimental spectrum was fitted to the Helfrich model for membrane fluctuations (43):

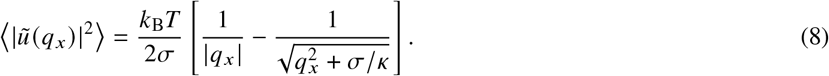

where a correction was incorporated for the finite camera integration time *τ*, which attenuates measured fluctuations at high wave vectors due to temporal averaging (50). The mode relaxation time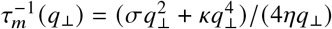, where *η* is the solvent viscosity, determines the extent of this attenuation. The bending rigidity *κ* and membrane tension *σ* were determined via nonlinear least-squares fitting using the Levenberg-Marquardt algorithm, with optimization performed in logarithmic space to ensure uniform weighting across the multi-decade range of fluctuation amplitudes. Following Pécréaux *et al*. (49), the first five modes were excluded due to spherical geometry effects (51), and high-*q* modes with ⟨|*ũ*|^2^⟩ < 10^−22^ m^3^ were excluded because they were below the detection noise floor.

To determine whether membrane fluctuations deviate from simple thermal relaxation under responsive conditions, the autocorrelation function (ACF) of each mode was computed as ⟨h_*q*_ (*t*′) h_*q*_ (*t*′ + *t*)⟩ to characterize the temporal relaxation behavior of the membrane. Deviations from a single-exponential decay (characteristic of thermal relaxation) were analyzed using a double-exponential model (52):

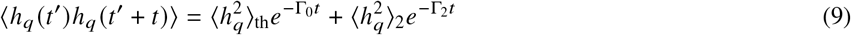

where Γ_0_ = (*σq κq*^3^) 4*η* represents passive relaxation and Γ_2_ corresponds to an additional relaxation process that deviate from simple thermal-equilibrium behavior, possibly arising from charge redistribution or IL protonation, or from ATP binding to ATPL.

### Micropipette aspiration

To assess the apparent membrane area expansion modulus (*K*_app_) of GUVs, micropipette aspiration studies were performed following established protocols (53–55). Custom borosilicate glass capillaries with minimal variation in wall thickness were obtained from Sutter Instrument (Novato, CA). Tips were incubated first in a 1% bovine serum albumin (BSA) coating solution for 5-10 minutes and then in Milli-Q water before each experiment. Micropipettes with a diameter of 8-10 *µ*m were used to aspirate GUVs, which have a diameter in the range of 10-15 *µ*m. Vesicles were partially aspirated into the micropipette with a vacuum pump (Welch Vacuum, Monroe, LA), which has a maximum pressure of 70 mbar. The aspiration pressure was precisely controlled using a digital pressure-based flow controller (Flow EZ−, Fluigent, Le Kremlin-Bicêtre, France) and a micromanipulator (MPC-200 and R0E-200, Sutter Instrument, Novato, CA). Micropipette aspiration experiments were conducted on the same day as GUV preparation.

Prepared GUVs were transferred to an observation chamber mounted on an inverted confocal laser scanning microscope (CLSM) (Carl Zeiss Microimaging GmbH, Germany) and visualized using bright-field imaging. The relative area dilation (Δ*A*/*A*) was obtained from changes in aspirated length, *L*_*p*_, and plotted against the applied membrane tension Σ = (Δ*P R*_*p*_) / (2 (1 − *R*_*p*_/*R*_*v*_)), where *R*_*p*_ is the pipette radius and *R*_*v*_ is the vesicle radius. From the linear regime of the Δ*A*/*A* vs Σ curve, the apparent area expansion modulus *K*_app_ was determined. For large deformations, thermal fluctuations were suppressed, allowing determination of the bending rigidity *κ* by fitting fluctuation spectra before and after aspiration (not shown). Vesicles showing visible membrane defects were excluded from the measurements because such defects could artificially alter the *K*_app_ values due to abnormal stretching. Image analysis was performed using ImageJ software (56) to measure pipette diameter, vesicle diameter, and the projection length of the vesicle inside the pipette.

## RESULTS AND DISCUSSION

### Lipid spontaneous curvature determined by SAXS

The elastic properties of membranes are strongly influenced by their spontaneous curvature *J*_0_, as described by the Helfrich elastic formalism. In this framework, the membrane bending energy depends on the deviation of the total curvature *J* = *C*_1_ + *C*_2_ from the spontaneous curvature, as well as on the Gaussian curvature *K* = *C*_1_*C*_2_. The energetic contributions associated with these deformations are weighted by the bending modulus *κ* and the Gaussian modulus 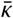, respectively (43, 44).

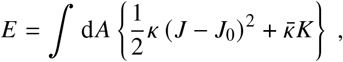

To evaluate how the incorporation of ILs and ATPL modulates the elastic properties of EEM, we determined the spontaneous curvature *J*_0_ of a judiciously chosen lipid mixture from SAXS measurements of inverse hexagonal *H*_II_ phases, following the geometrical analysis described by Gillams *et al*. (42). Here, DOPE lipids self-assembled into an *H*_∥_ phase were doped with 10 mol % of the lipids of interest. This approach preserves the integrity of the *H*_∥_ host matrix while taking advantage of the fact that its lattice parameter *a* is highly sensitive to the presence of trace amounts of additional lipids. The lattice parameter *a* was measured at different temperatures (25 to 70 °C) under steady-state and responsive conditions, i.e., in NaOAc buffer at pH 5.0-5.5 that protonates ILs and 1 mM ATP that binds to ATPL headgroups.

The results in Figure 2A clearly show a linear relationship consistent with Equation 4 with slope 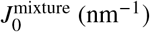, which represents the curvature elastic energy as a function of temperature. The *J*_0_ values determined by SAXS shown in Figure 2B suggest that incorporation of ILs/ATPL in the EEM induces a clear shift of *J*_0_ towards more negative values relative to pure EEMs, with the magnitude of the shift increasing under inducing conditions of low pH (ILs) and ATP addition (ATPL).

**Figure 2:**
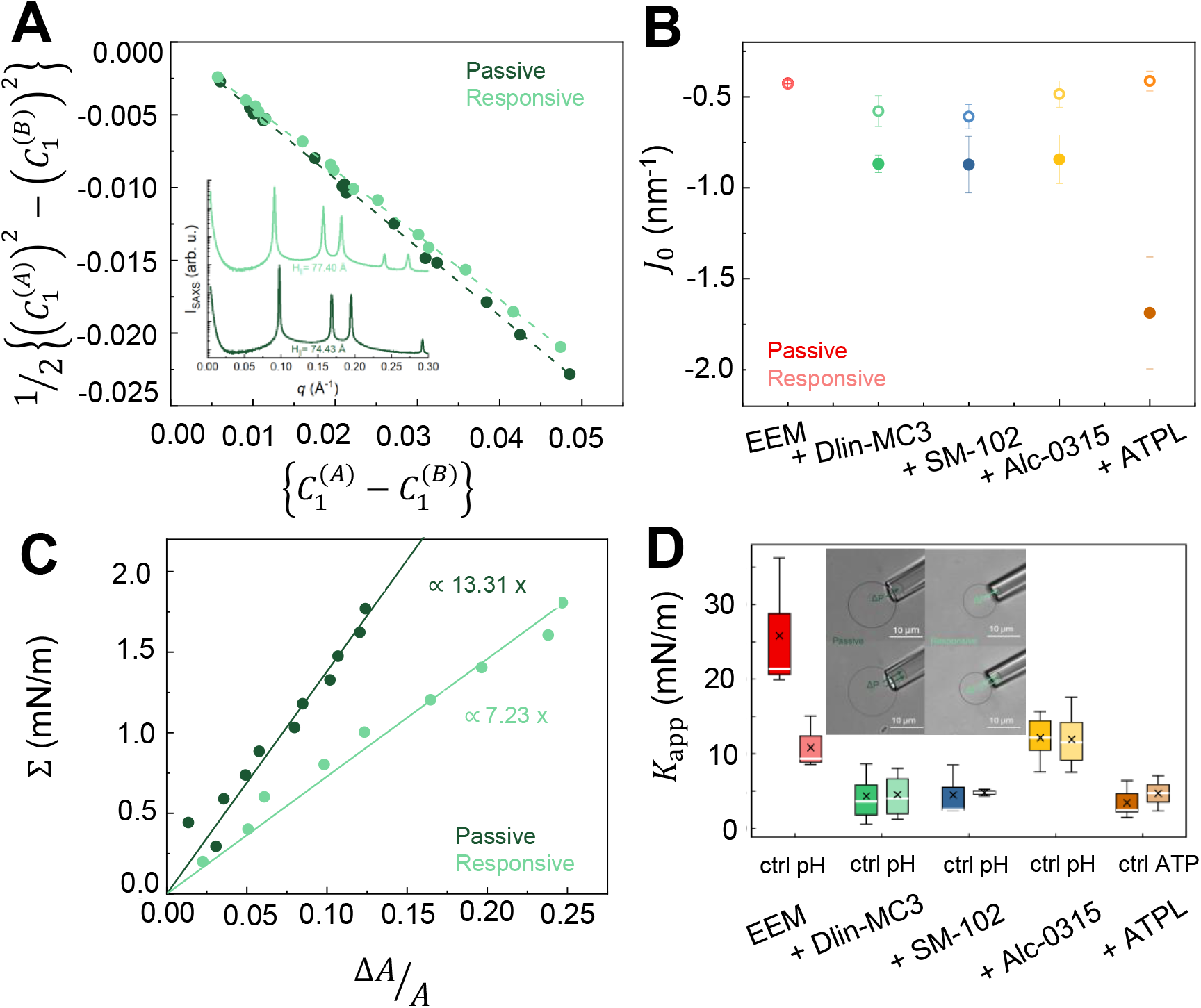
Spontaneous curvature and membrane deformability induced by ILs/ATPL. (A) Linearized analysis of Eq. 4 for DOPE + EEM with Dlin-MC3. The linear fits yield *J*_0_ of the mixture as the slope. The inset shows the SAXS profile of the same sample at 25 °C. (B) Extracted *J*_0_ of EEM membranes with different ILs and ATPL under steady-state and responsive conditions. (C,D) Micropipette-aspiration results for giant unilamellar vesicles (GUVs). (C) Representative tension (Σ) vs. fractional change in surface area for a single GUV containing the ionizable lipid Dlin-MC3 under steady-state and responsive conditions. The slope of the linear fit yields the apparent area expansion modulus *K*_app_. (D) *K*_app_ values of GUVs containing different responsive lipids (color-coded as in panel B) under steady-state and responsive conditions. The white line and black cross indicate the median and mean values, respectively. The inset shows 2D cross-sectional bright-field images of representative Dlin-MC3 GUVs used for micropipette aspiration. The top image shows the vesicle prior to aspiration; the bottom image shows the change in aspirated membrane length induced by applied pressure.

A more negative *J*_0_ signifies that the membrane’s elastic energy minimum moves toward inverted curvature geometries, such as those associated with stalk intermediates, fusion pores, and other nonlamellar deformations (57, 58). Physically, this indicates that IL/ATPL incorporation biases the membrane toward curved states and lowers the energetic cost of high-curvature membrane deformations. The results for ATPL can be rationalized as follows: as ATP binds to zinc ions in the headgroup, the alkyl tails splay and the lipid shape changes towards inverted geometries. This has been putatively linked to the benefits of formulating ATPL into liposomal delivery systems (38).

The behavior of ILs, however, is less intuitive. Upon protonation, IL headgroups become positively charged, and the headgroup-to-headgroup electrostatic repulsion would be expected to effectively increase the cross-sectional area per headgroup, thereby reducing the molecular packing parameter and making the lipid appear less inverted (59). However, clinically relevant ILs, especially SM-102 and Alc-0315, possess exceptionally bulky, highly branched hydrophobic tails and remain strongly cone-shaped molecules. Consequently, the increase in effective headgroup area associated with protonation is expected to represent only a relatively small perturbation compared with the large hydrophobic volume of the lipid. Moreover, in the present study, ILs are present only as minor components dispersed within an EEM mimic composed primarily of *H*_∥_ host matrix. Under these conditions, protonation-induced electrostatic repulsion is more likely to promote lateral separation of individual IL molecules rather than collective packing. This separation is expected to redistribute the lateral stress profile within the membrane, while the bulky hydrophobic tails couple to neighboring curvature-preferring lipids such as DOPE, collectively amplifying the membrane’s tendency toward negative spontaneous curvature. These observations emphasize that the molecular shape alone is insufficient to explain the membrane response to IL/ATPL incorporation. Although packing parameter arguments provide useful intuition at the molecular level, curvature and elastic moduli are emergent continuum properties that arise from the collective organization of many lipids within the membrane.

Indeed, although the SAXS *J*_0_ measurements were conducted using an *H*_∥_ host matrix, endosomal membranes are constrained to adopt an overall lamellar geometry, leading to monolayers not being able to fully adopt their preferred curvature. This mismatch generates curvature frustration and a corresponding elastic energy penalty that can be approximated by *F* ≈ 1/2*κ* (*J* − *J*_0_)^2^ (60, 61). As *J*_0_ becomes more negative while the local mean curvature *J ≈* 0, this curvature frustration energy increases. This stored elastic energy may be partially relieved through enhanced membrane undulations and increased curvature fluctuations, which can manifest experimentally as a reduction in the apparent bending rigidity. Understanding how new lipid chemistries influence membrane mechanics requires consideration of both local molecular packing and the global elastic response of the membrane. In the following sections, we evaluate how the continuum mechanical properties of GUV-based early endosomal membrane mimics respond to the inclusion of lipids triggered by pH changes and/or ATP fluxes.

### Area expansion from micropipette aspiration

Micropipette aspiration (MPA) is a well-established method for measuring the elastic behavior of model (53, 54, 62–65) and biological membranes (54, 66, 67). By measuring membrane deformations in response to an applied suction pressure, MPA enables quantitative comparison of membrane tension, apparent area expansion modulus *K*_app_, and membrane viscosity under different experimental conditions. Although MPA does not directly measure the intrinsic membrane stretch modulus, it provides a robust framework for comparing relative mechanical changes between samples. In particular, *K*_app_ quantifies the membrane’s resistance to in-plane area expansion and, at low membrane tensions, reflects contributions from both elastic membrane stretching and suppression of thermal undulations. Thus, changes in *K*_app_, associated with the incorporation of ILs/ATPL or with exposure to environmental stimuli (e.g., acidic pH or ATP binding) provide a sensitive measure of changes in membrane mechanical behavior as a function of membrane composition and environmental conditions.

Figures 2C and D show representative aspiration results and the *K*_app_ distributions, respectively, for five membrane compositions containing three different ILs and ATPL. Figure 2C plots tension Σ versus fractional change in surface area for a representative GUV in both steady-state and triggering conditions. The change in slope indicates the reduced resistance of the membrane containing ILs under low pH conditions to expansion. Figure 2D shows the *K*_app_ distributions for GUVs comprising different responsive lipids under steady-state and triggering conditions; white lines and black crosses indicate the medians and means, respectively. The insets illustrate the two-dimensional (2D) cross-sectional bright-field images of Dlin-MC3-based GUVs captured at the initial suction pressure and after subsequent increases in pressure, Δ*P*. Suction was applied until either the vesicles were fully pulled in or sufficient data points were collected. Under steady-state conditions, the relatively stiffer vesicles allowed the acquisition of more data points, whereas under responsive conditions, membrane rupture or complete aspiration occurred more frequently. For each membrane composition, the number of vesicles analyzed was greater than five (*n* > 5).

Membranes containing Dlin-MC3, SM-102, Alc-0315, and ATPL exhibited softening relative to membranes with the pure EEM composition. Although the mean values appear slightly higher, the box plots indicate that the changes lie within the margin of error, and the overall trend is a decrease in effective membrane stiffness. This apparent discrepancy can be attributed to the nature of the MPA technique, which is sensitive to factors such as vesicle–pipette interactions, pressure fluctuations, and variations in the robotic manipulation of the vesicles as the medium is replaced by ATP or buffer solutions. Nevertheless, it remains a highly effective comparative method, complementary to the fluctuation analysis and SAXS measurements discussed in separate sections. The observed decrease in *K*_app_ values indicates that incorporation of ILs and ATPL reduces the membrane’s resistance to area expansion, making the membrane more readily deformable under applied tension. This reduction can be attributed to changes in headgroup chemistry, variations in *J*_0_, and enhanced headgroup–headgroup repulsion, which promote negative spontaneous curvature and relaxation of membrane tension, as discussed above.

### pH- and ATP-dependent enhancement of membrane fluctuations

To evaluate alterations in membrane stiffness caused by the incorporation of ILs/ATPL into EEM mimetics without the complication of pipette interactions, we characterized membrane stiffness by vesicle fluctuation analysis under steady-state and responsive conditions. Figure 3 shows representative results for EEM GUVs with 10 mol% Dlin-MC3.

**Figure 3:**
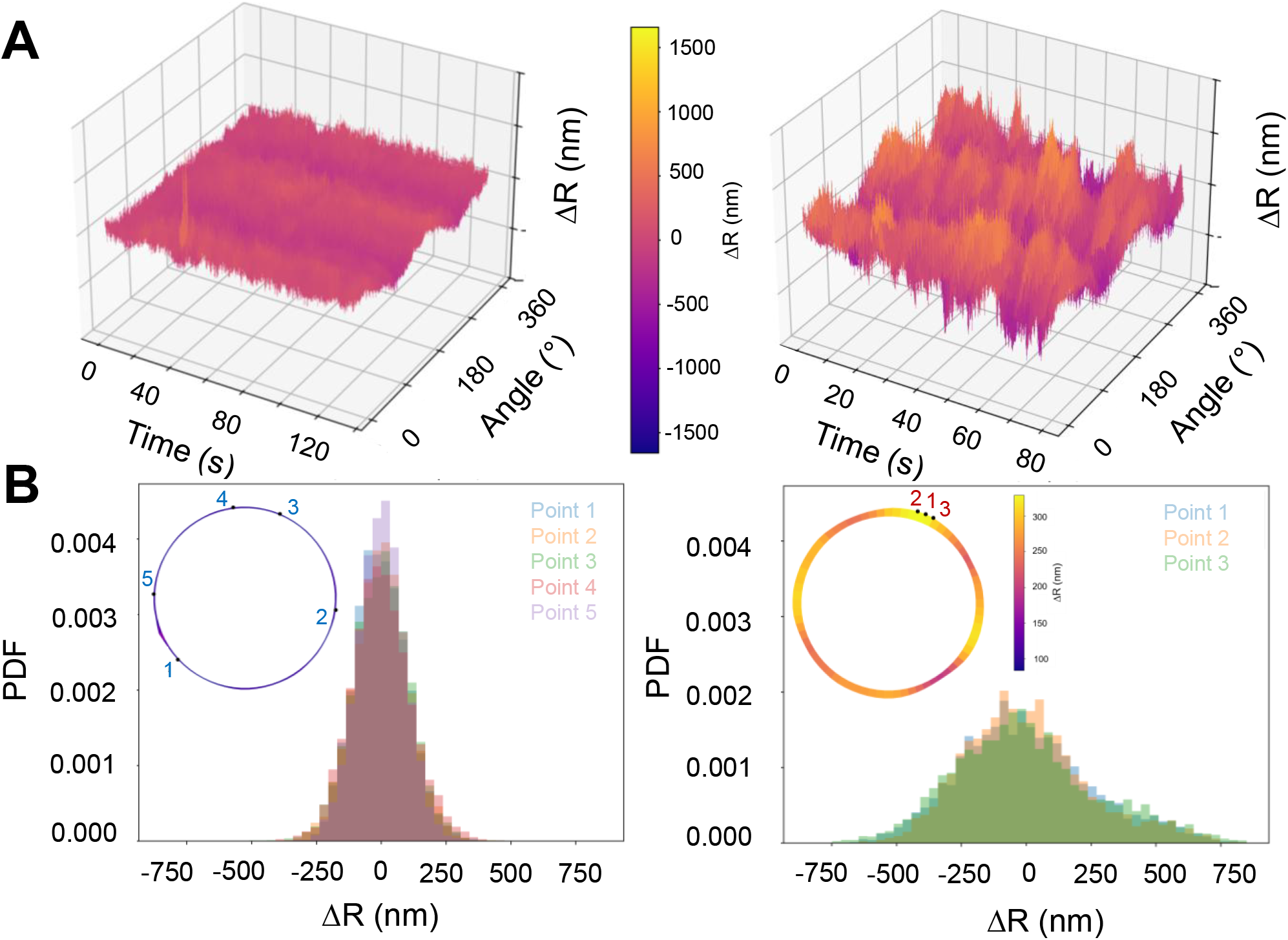
Enhanced membrane fluctuations of a single EEM + Dlin-MC3 vesicle under steady-state and responsive conditions. (A) Spatiotemporal maps of radial displacements, Δ*R* (*θ, t*), as a function of angle and time. Passive membranes (left) display low-amplitude, relatively homogeneous fluctuations, whereas responsive membranes (right) show larger-amplitude, spatially heterogeneous fluctuations with pronounced temporal variability. (B) Probability distribution functions (PDFs) of local radial displacements, Δ*R*, sampled at multiple contour positions. Steady-state membranes exhibit narrow, symmetric distributions, while responsive membranes show broader distributions with increased variance. Insets show the time-averaged fluctuation amplitude maps of the vesicle contour and the spatial distribution of radial displacement magnitude Δ*R* (nm). Under steady-state conditions, fluctuations are low and relatively uniform along the contour. In contrast, responsive membranes exhibit higher fluctuation amplitudes with localized regions of enhanced activity.

The insets in Figure 3B show the time-averaged fluctuation amplitude maps of membranes under steady-state (left) and responsive conditions (after adding NaOAc buffer, right). The fluctuation amplitude maps clearly show that under steady-state conditions, the fluctuation intensity remains low and spatially uniform along the contour, consistent with passive, thermally-driven behavior. In contrast, after addition of low pH buffer, fluctuations become more pronounced and distinct regions with high fluctuation amplitude emerge, appearing as bright domains in the map. These regions correspond to transient, localized shape changes, including small protrusions and indentations, that vary dynamically with time. Figure 3A shows the three-dimensional (3D) spatiotemporal representation of radial displacements, Δ*R* (*θ, t*), revealing how the membrane shape evolves (Supporting Information). For the control vesicles, the surface remains mostly flat, exhibiting small, uncorrelated fluctuations around the mean contour. In acidic conditions, however, the amplitude and frequency of fluctuations increase dramatically, producing dynamic, wave-like patterns with excursions exceeding several hundred nanometers. This enhancement of fluctuation amplitude provides clear visual evidence that the responsiveness of ILs to acidic pH introduces enhanced membrane fluctuations, consistent with a deviation from thermal equilibrium.

To quantify these deviations, we analyzed the probability distribution function (PDF) of local height fluctuations, h (*t*) − ⟨h⟩. Figure 3B shows the PDFs for the marked points in its insets. At equilibrium, the fluctuations follow near-Gaussian distributions, as expected from thermally driven membranes. Consistent with this observation, steady-state membranes exhibit narrow, symmetric PDFs across all sampled contour regions. However, under acidic conditions, the PDFs display noticeable deviations from Gaussianity, particularly at fluctuation “hot spots” where asymmetric, heavy-tailed distributions emerge. These features reflect intermittent, large-amplitude events that deviate from the near-Gaussian fluctuation behavior observed under steady-state conditions. At a global level, combining the fluctuations across the entire contour further highlights this transition. Passive membranes display a single, narrow Gaussian distribution, whereas responsive membranes show substantial broadening, indicating an overall increase in fluctuation amplitude (Supporting Information). The emergence of non-Gaussian PDFs under acidic or ATP conditions, therefore, provides experimental evidence that protonation of IL headgroups and ATP binding are associated with enhanced, spatially heterogeneous membrane fluctuations. The softening effect, manifested as broader, asymmetric PDFs, precedes and quantitatively predicts the reduction in bending rigidity and surface tension described in the following section.

### Relaxation dynamics and membrane stiffness

To quantify the dynamics and mechanical response of these membranes, we analyzed both the fluctuation spectra and temporal relaxation behavior of the contour modes (Figure 4). The fluctuation spectrum for a representative vesicle is shown in Figure 4A. Under steady-state conditions, the spectrum follows the expected Helfrich scaling, with fluctuation amplitudes decreasing with increasing wave vector. In contrast, under responsive conditions, the spectrum is elevated across all modes, indicating enhanced fluctuations consistent with membrane softening. Fits to the Helfrich model (Eq. 8) were used to determine the bending rigidity *κ*. The experimentally determined values of bending rigidity for different ILs are summarized in Figure 4B. Under steady-state conditions, membranes exhibit *κ* values in the range of ≈ 15-20 *k*_*B*_*T*, consistent with fluid lipid bilayers containing unsaturated lipids and cholesterol (68). Upon addition of low pH buffer or ATP, *κ* decreases systematically across all compositions, indicating a softening of the membrane. This reduction reflects an increased susceptibility to bending deformations, consistent with the enhanced fluctuations observed in the spectra.

**Figure 4:**
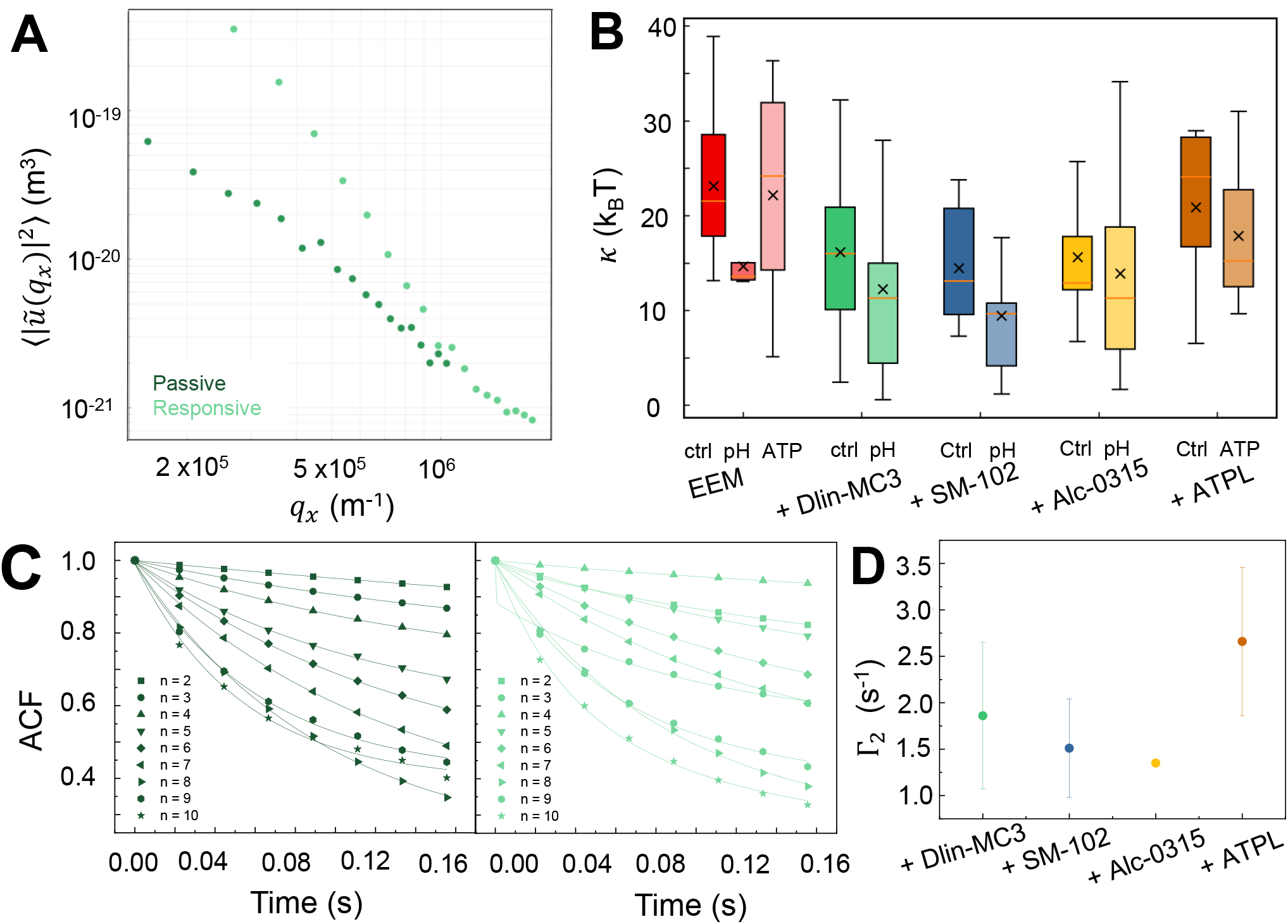
Fluctuation spectra a nd relaxation dynamics of membranes under steady-state and responsive conditions. (A) Fluctuation spectrum ⟨|*ũ* (*q*_*x*_) |^2^⟩ for a representative EEM + Dlin-MC3 vesicle as a function of wave vector *q*. Fluctuation amplitudes decrease with increasing *q*, consistent with membrane undulation behavior. (B) Experimentally determined bending rigidity *κ* from fluctuation spectra for different membrane compositions under steady-state and responsive conditions. Distributions are shown for pure EEM membranes and EEM membranes containing Dlin-MC3, SM-102, Alc-0315, or ATPL. The orange line and black cross indicate the median and mean values, respectively. (C) Autocorrelation functions (ACFs) of different contour modes (*n* = 2–10) under steady-state (left) and responsive (right) conditions for the same vesicle containing EEM+Dlin-MC3, showing the temporal decay of membrane fluctuations. Solid lines represent fits to a single-exponential function for passive conditions and a double-exponential function for responsive conditions. (D) Characteristic slow relaxation rates Γ_2_ extracted from ACF analysis for different lipid compositions under responsive conditions.

To characterize the temporal nature of these fluctuations, we determined the ACFs of contour modes (Figure 4C). Under steady-state conditions, the ACFs exhibit a relatively smooth, approximately single-exponential decay, consistent with thermally driven relaxation. In contrast, responsive membranes show a markedly slower and more heterogeneous decay, with clear deviations from single-exponential behavior. These features indicate the presence of additional relaxation processes beyond equilibrium thermal fluctuations. Notably, the responsive ACFs also exhibit a different and non-monotonic ordering with mode number, indicating that the relaxation dynamics are altered across spatial modes. Rather than following a single relaxation process governed by membrane elasticity and viscous damping, the dynamics under responsive conditions exhibit multiple relaxation components. By fitting the decays to a double-exponential function (Eq. 9), the additional relaxation rate Γ_2_ can be determined to characterize the slower relaxation process observed under responsive conditions (Figure 4D). This treatment is consistent with previous descriptions of nonequilibrium membrane fluctuations, in which an additional relaxation mode accompanies the conventional thermal relaxation process (69). Mechanistically, these effects may arise from chemical changes within the membrane. In IL-containing membranes, protonation of the headgroups under acidic conditions may alter lipid packing and generate curvature stress. Similarly, in ATPL-containing vesicles, ATP binding may alter molecular packing and preferred curvature. In both cases, these changes are associated with enhanced fluctuations, reduced effective bending rigidity, and slower relaxation, indicating membrane dynamics that deviate from simple thermal-equilibrium behavior.

## CONCLUSION

Our results provide a new quantitative understanding of how ILs/ATPL used in state-of-the-art LNP delivery systems may impact the biophysical properties of endosomal membranes during a cellular delivery event. By combining SAXS, micropipette aspiration, and vesicle fluctuation analysis, we conducted a comprehensive study showing that protonation or ATP binding is associated with membrane softening and enhanced, non-Gaussian fluctuations that deviate from simple thermal-equilibrium behavior. These observations are consistent with a shift in lipid spontaneous curvature towards more negative values, which introduces curvature frustration in quasi-spherical vesicles. Rather than inducing immediate large-scale shape transformations, this stored elastic stress is dissipated through amplified, spatially heterogeneous fluctuations. Importantly, this behavior is observed across both IL- and ATPL-containing systems, suggesting a common physical mechanism in which stimulus-induced changes in lipid packing and preferred curvature couple to membrane bending and fluctuation dynamics. Such membrane remodeling events provide a plausible pathway for transient destabilization of early endosomal membranes, facilitating escape. Broadly, this work establishes a quantitative framework linking lipid molecular properties to membrane mechanics and dynamics, offering design principles for next-generation delivery systems that exploit controlled curvature stress and enhanced fluctuations to improve intracellular transport.

## AUTHOR CONTRIBUTIONS

T.K. and C.L. designed the study. T.K. carried out the SAXS experiments and analyzed the data. T.K. and B.S.S. carried out micropipette aspiration and analyzed the data. Y.L. carried out vesicle fluctuation experiments, and T.K. and Y.L. analyzed the data. C.S. oversaw the fluctuation data analysis. M.B.M., J.L., and M.B. synthesized and provided the ATP-responsive lipids. T.K. and C.L. wrote the article with input from Y.L., B.S.S., C.S., M.B.M., J.L., and M.B.

## ACKNOWLEDGMENTS

This work was supported by the NIH under Grant No. R01GM143723-01A1. This research used resources of the Advanced Photon Source, beamline 12-ID-B, a US Department of Energy (DOE) Office of Science User Facility operated for the DOE Office of Science by Argonne National Laboratory under Contract No. DE-AC02-06CH11357. Confocal microscopy images were acquired using an instrument funded by the Office of Naval Research (ONR) DURIP–Defense University Research Instrumentation Program, Grant No. N000141812087. This research was conducted in part in the Materials Research Laboratory at the University of Illinois Urbana-Champaign. We thank Max Baliga for help with in-house SAXS measurements, Puquan Pan and James Tallman for helping us develop the contour-tracking and analysis code.

## REFERENCES

1. Cobb, M., 2015. Who discovered messenger RNA? Current Biology 25:R526–R532.

2. Sahin, U., K. Karikó, and Ö. Türeci, 2014. mRNA-based therapeutics—developing a new class of drugs. Nature reviews Drug discovery 13:759–780.

3. Wolff, J. A., R. W. Malone, P. Williams, W. Chong, G. Acsadi, A. Jani, and P. L. Felgner, 1990. Direct gene transfer into mouse muscle in vivo. Science 247:1465–1468.

4. Karikó, K., M. Buckstein, H. Ni, and D. Weissman, 2005. Suppression of RNA recognition by Toll-like receptors: the impact of nucleoside modification and the evolutionary origin of RNA. Immunity 23:165–175.

5. Karikó, K., H. Muramatsu, F. A. Welsh, J. Ludwig, H. Kato, S. Akira, and D. Weissman, 2008. Incorporation of pseudouridine into mRNA yields superior nonimmunogenic vector with increased translational capacity and biological stability. Molecular therapy 16:1833–1840.

6. Hou, X., T. Zaks, R. Langer, and Y. Dong, 2021. Lipid nanoparticles for mRNA delivery. Nature Reviews Materials 6:1078–1094.

7. Tenchov, R., R. Bird, A. E. Curtze, and Q. Zhou, 2021. Lipid nanoparticles from liposomes to mRNA vaccine delivery, a landscape of research diversity and advancement. ACS nano 15:16982–17015.

8. Anderson, E. J., N. G. Rouphael, A. T. Widge, L. A. Jackson, P. C. Roberts, M. Makhene, J. D. Chappell, M. R. Denison, L. J. Stevens, A. J. Pruijssers, et al., 2020. Safety and immunogenicity of SARS-CoV-2 mRNA-1273 vaccine in older adults. New England Journal of Medicine 383:2427–2438.

9. Baden, L. R., H. M. El Sahly, B. Essink, K. Kotloff, S. Frey, R. Novak, D. Diemert, S. A. Spector, N. Rouphael, C. B. Creech, et al., 2021. Efficacy and safety of the mRNA-1273 SARS-CoV-2 vaccine. New England journal of medicine 384:403–416.

10. Polack, F. P., S. J. Thomas, N. Kitchin, J. Absalon, A. Gurtman, S. Lockhart, J. L. Perez, G. Pérez Marc, E. D. Moreira, C. Zerbini, et al., 2020. Safety and efficacy of the BNT162b2 mRNA Covid-19 vaccine. New England journal of medicine 383:2603–2615.

11. Akinc, A., M. A. Maier, M. Manoharan, K. Fitzgerald, M. Jayaraman, S. Barros, S. Ansell, X. Du, M. J. Hope, T. D. Madden, et al., 2019. The Onpattro story and the clinical translation of nanomedicines containing nucleic acid-based drugs. Nature nanotechnology 14:1084–1087.

12. Sahay, G., W. Querbes, C. Alabi, A. Eltoukhy, S. Sarkar, C. Zurenko, E. Karagiannis, K. Love, D. Chen, R. Zoncu, et al., 2013. Efficiency of siRNA delivery by lipid nanoparticles is limited by endocytic recycling. Nature biotechnology 31:653–658.

13. Wittrup, A., A. Ai, X. Liu, P. Hamar, R. Trifonova, K. Charisse, M. Manoharan, T. Kirchhausen, and J. Lieberman, 2015. Visualizing lipid-formulated siRNA release from endosomes and target gene knockdown. Nature biotechnology 33:870–876.

14. Zheng, L., S. R. Bandara, Z. Tan, and C. Leal, 2023. Lipid nanoparticle topology regulates endosomal escape and delivery of RNA to the cytoplasm. Proceedings of the National Academy of Sciences 120:e2301067120. https://pnas.org/doi/10.1073/pnas.2301067120.

15. Zheng, L., M. Baliga, S. F. Gallagher, A. Gao, J. Rueben, Y. K. Go, M. Deserno, and C. Leal, 2026. Introducing a fusogenicity metric for lipid nanoparticle formulation. bioRxiv 2026–03.

16. Degors, I. M., C. Wang, Z. U. Rehman, and I. S. Zuhorn, 2019. Carriers break barriers in drug delivery: endocytosis and endosomal escape of gene delivery vectors. Accounts of chemical research 52:1750–1760.

17. Patel, S., J. Kim, M. Herrera, A. Mukherjee, A. V. Kabanov, and G. Sahay, 2019. Brief update on endocytosis of nanomedicines. Advanced drug delivery reviews 144:90–111.

18. Gilleron, J., W. Querbes, A. Zeigerer, A. Borodovsky, G. Marsico, U. Schubert, K. Manygoats, S. Seifert, C. Andree, M. Stöter, et al., 2013. Image-based analysis of lipid nanoparticle–mediated siRNA delivery, intracellular trafficking and endosomal escape. Nature biotechnology 31:638–646.

19. Kauffman, K. J., J. R. Dorkin, J. H. Yang, M. W. Heartlein, F. DeRosa, F. F. Mir, O. S. Fenton, and D. G. Anderson, 2015. Optimization of Lipid Nanoparticle Formulations for mRNA Delivery in Vivo with Fractional Factorial and Definitive Screening Designs. Nano Letters 15:7300–7306. https://pubs.acs.org/doi/10.1021/acs.nanolett.5b02497.

20. Patel, S., N. Ashwanikumar, E. Robinson, A. DuRoss, C. Sun, K. E. Murphy-Benenato, C. Mihai, O. Almarsson, and G. Sahay, 2017. Boosting Intracellular Delivery of Lipid Nanoparticle-Encapsulated mRNA. Nano Letters 17:5711–5718.10.1021/acs.nanolett.7b02664, publisher: American Chemical Society.

21. Patel, S., N. Ashwanikumar, E. Robinson, Y. Xia, C. Mihai, J. P. Griffith, S. Hou, A. A. Esposito, T. Ketova, K. Welsher, J. L. Joyal, O. Almarsson, and G. Sahay, 2020. Naturally–occurring cholesterol analogues in lipid nanoparticles induce polymorphic shape and enhance intracellular delivery of mRNA. Nature Communications 11:983. https://www.nature.com/articles/s41467-020-14527-2, publisher: Nature Publishing Group.

22. Semple, S. C., A. Akinc, J. Chen, A. P. Sandhu, B. L. Mui, C. K. Cho, D. W. Y. Sah, D. Stebbing, E. J. Crosley, E. Yaworski, I. M. Hafez, J. R. Dorkin, J. Qin, K. Lam, K. G. Rajeev, K. F. Wong, L. B. Jeffs, L. Nechev, M. L. Eisenhardt, M. Jayaraman, M. Kazem, M. A. Maier, M. Srinivasulu, M. J. Weinstein, Q. Chen, R. Alvarez, S. A. Barros, S. De, S. K. Klimuk, T. Borland, V. Kosovrasti, W. L. Cantley, Y. K. Tam, M. Manoharan, M. A. Ciufolini, M. A. Tracy, A. de Fougerolles, I. MacLachlan, P. R. Cullis, T. D. Madden, and M. J. Hope, 2010. Rational design of cationic lipids for siRNA delivery. Nature Biotechnology 28:172–176. https://www.nature.com/articles/nbt.1602, publisher: Nature Publishing Group.

23. Barriga, H. M. G., M. N. Holme, and M. M. Stevens, 2019. Cubosomes: The Next Generation of Smart Lipid Nanoparticles? Angewandte Chemie International Edition 58:2958–2978. https://onlinelibrary.wiley.com/doi/10.1002/anie.201804067.

24. Leung, S. S., and C. Leal, 2019. The stabilization of primitive bicontinuous cubic phases with tunable swelling over a wide composition range. Soft matter 15:1269–1277.

25. Kim, H., Z. Song, and C. Leal, 2017. Super-swelled lyotropic single crystals. Proceedings of the National Academy of Sciences 114:10834–10839.

26. Kang, M., H. Kim, and C. Leal, 2016. Self-organization of nucleic acids in lipid constructs. Current opinion in colloid & interface science 26:58–65.

27. Rueben, J., A. Jayaraman, M. K. Mahanthappa, and C. Leal, 2022. Near-infrared-triggered reversible transformations of gold nanorod-laden lipid assemblies: Implications for cellular delivery. ACS applied nano materials 5:710–717.

28. Porras-Gomez, M., and C. Leal, 2019. Lipid-based liquid crystalline films and solutions for the delivery of cargo to cells. Liquid crystals reviews 7:167–182.

29. Kim, H., and C. Leal, 2015. Cuboplexes: Topologically active siRNA delivery. ACS nano 9:10214–10226.

30. Leal, C., N. F. Bouxsein, K. K. Ewert, and C. R. Safinya, 2010. Highly efficient gene silencing activity of siRNA embedded in a nanostructured gyroid cubic lipid matrix. Journal of the American Chemical Society 132:16841–16847.

31. Leal, C., K. K. Ewert, R. S. Shirazi, N. F. Bouxsein, and C. R. Safinya, 2011. Nanogyroids incorporating multivalent lipids: enhanced membrane charge density and pore forming ability for gene silencing. Langmuir 27:7691–7697.

32. Omo-Lamai, S., Y. Wang, M. N. Patel, A. Milosavljevic, D. Zuschlag, S. Poddar, J. Wu, L. Wang, F. Dong, C. Espy, et al., 2025. Limiting endosomal damage sensing reduces inflammation triggered by lipid nanoparticle endosomal escape. Nature nanotechnology 20:1285–1297.

33. Lee, Y., M. Jeong, J. Park, H. Jung, and H. Lee, 2023. Immunogenicity of lipid nanoparticles and its impact on the efficacy of mRNA vaccines and therapeutics. Experimental & Molecular Medicine 55:2085–2096.

34. Chaudhary, N., L. N. Kasiewicz, A. N. Newby, M. L. Arral, S. S. Yerneni, J. R. Melamed, S. T. LoPresti, K. C. Fein, D. M. Strelkova Petersen, S. Kumar, et al., 2024. Amine headgroups in ionizable lipids drive immune responses to lipid nanoparticles by binding to the receptors TLR4 and CD1d. Nature Biomedical Engineering 8:1483–1498.

35. Han, X., H. Zhang, K. Butowska, K. L. Swingle, M.-G. Alameh, D. Weissman, and M. J. Mitchell, 2021. An ionizable lipid toolbox for RNA delivery. Nature communications 12:7233.

36. Paramasivam, P., C. Franke, M. Stöter, A. Höijer, S. Bartesaghi, A. Sabirsh, L. Lindfors, M. Y. Arteta, A. Dahlén, A. Bak, S. Andersson, Y. Kalaidzidis, M. Bickle, and M. Zerial, 2021. Endosomal escape of delivered mRNA from endosomal recycling tubules visualized at the nanoscale. Journal of Cell Biology 221:e202110137. 10.1083/jcb.202110137.

37. Sayers, E. J., S. E. Peel, A. Schantz, R. M. England, M. Beano, S. M. Bates, A. S. Desai, S. Puri, M. B. Ashford, and A. T. Jones, 2019. Endocytic Profiling of Cancer Cell Models Reveals Critical Factors Influencing LNP-Mediated mRNA Delivery and Protein Expression. Molecular Therapy 27:1950–1962. https://www.cell.com/molecular-therapy-family/molecular-therapy/abstract/S1525-0016(19)30357-0, publisher: Elsevier.

38. Lou, J., J. A. Schuster, F. N. Barrera, and M. D. Best, 2022. ATP-responsive liposomes via screening of lipid switches designed to undergo conformational changes upon binding phosphorylated metabolites. Journal of the American Chemical Society 144:3746–3756.

39. Mathivet, L., S. Cribier, and P. F. Devaux, 1996. Shape change and physical properties of giant phospholipid vesicles prepared in the presence of an AC electric field. Biophysical journal 70:1112–1121.

40. Kobayashi, T., M.-H. Beuchat, J. Chevallier, A. Makino, N. Mayran, J.-M. Escola, C. Lebrand, P. Cosson, T. Kobayashi, and J. Gruenberg, 2002. Separation and characterization of late endosomal membrane domains. Journal of biological chemistry 277:32157–32164.

41. Bergstrand, N., M. C. Arfvidsson, J.-M. Kim, D. H. Thompson, and K. Edwards, 2003. Interactions between pH-sensitive liposomes and model membranes. Biophysical chemistry 104:361–379.

42. Gillams, R. J., T. Nylander, T. S. Plivelic, M. K. Dymond, and G. S. Attard, 2014. Formation of inverse topology lyotropic phases in dioleoylphosphatidylcholine/oleic acid and dioleoylphosphatidylethanolamine/oleic acid binary mixtures. Langmuir 30:3337–3344.

43. Helfrich, W., 1973. Elastic properties of lipid bilayers: theory and possible experiments. Zeitschrift für Naturforschung C 28:693–703.

44. Germain, S., 1821. Recherches sur la théorie des surfaces élastiques. Mme. Ve. Courcier.

45. Dymond, M. K., R. J. Gillams, D. J. Parker, J. Burrell, A. Labrador, T. Nylander, and G. S. Attard, 2016. Lipid spontaneous curvatures estimated from temperature-dependent changes in inverse hexagonal phase lattice parameters: effects of metal cations. Langmuir 32:10083–10092.

46. Dymond, M. K., 2021. Lipid monolayer spontaneous curvatures: A collection of published values. Chemistry and Physics of Lipids 239:105117.

47. Ravi, N., V. Gabeur, Y.-T. Hu, et al., 2024. SAM 2: Segment Anything in Images and Videos. arXiv preprint arXiv:2408.00714

48. Bradski, G., 2000. The OpenCV Library. Dr. Dobb’s Journal of Software Tools.

49. Pécréaux, J., H.-G. Döbereiner, J. Prost, J.-F. Joanny, and P. Bassereau, 2004. Refined contour analysis of giant unilamellar vesicles. European Physical Journal E 13:277–290.

50. Faucon, J.-F., M. D. Mitov, P. Méléard, I. Bivas, and P. Bothorel, 1989. Bending elasticity and thermal fluctuations of lipid membranes. Theoretical and experimental requirements. Journal de Physique 50:2389–2414.

51. Milner, S. T., and S. A. Safran, 1987. Dynamical fluctuations of droplet microemulsions and vesicles. Physical Review A 36:4371–4379.

52. Milner, S. T., and S. Safran, 1987. Dynamical fluctuations of droplet microemulsions and vesicles. Physical Review A 36:4371.

53. Evans, E., K. Ritchie, and R. Merkel, 1995. Sensitive force technique to probe molecular adhesion and structural linkages at biological interfaces. Biophysical journal 68:2580–2587.

54. Hochmuth, R. M., 2000. Micropipette aspiration of living cells. Journal of biomechanics 33:15–22.

55. Rawicz, W., B. Smith, T. McIntosh, S. Simon, and E. Evans, 2008. Elasticity, strength, and water permeability of bilayers that contain raft microdomain-forming lipids. Biophysical journal 94:4725–4736.

56. Schneider, C. A., W. S. Rasband, and K. W. Eliceiri, 2012. NIH Image to ImageJ: 25 years of image analysis. Nature Methods 9:671–675. 10.1038/nmeth.2089.

57. Fan, Z.-A., K.-Y. Tsang, S.-H. Chen, and Y.-F. Chen, 2016. Revisit the Correlation between the Elastic Mechanics and Fusion of Lipid Membranes. Scientific Reports 6:31470. 10.1038/srep31470.

58. May, S., 2002. Structure and Energy of Fusion Stalks: The Role of Membrane Edges. Biophysical Journal 83:2969–2980. 10.1016/S0006-3495(02)75303-4.

59. Israelachvili, J. N., 2011. Intermolecular and surface forces. Academic press.

60. Helfrich, W., 1973. Elastic properties of lipid bilayers: theory and possible experiments. Zeitschrift für Naturforschung c 28:693–703.

61. Lipowsky, R., 1991. The conformation of membranes. Nature 349:475–481.

62. Evans, E. A., R. Waugh, and L. Melnik, 1976. Elastic area compressibility modulus of red cell membrane. Biophysical journal 16:585–595.

63. Needham, D., and R. S. Nunn, 1990. Elastic deformation and failure of lipid bilayer membranes containing cholesterol. Biophysical journal 58:997–1009.

64. Ratanabanangkoon, P., M. Gropper, R. Merkel, E. Sackmann, and A. P. Gast, 2003. Mechanics of streptavidin-coated giant lipid bilayer vesicles: A micropipet study. Langmuir 19:1054–1062.

65. Henriksen, J. R., and J. H. Ipsen, 2004. Measurement of membrane elasticity by micro-pipette aspiration. The European physical journal E 14:149–167.

66. González-Bermúdez, B., G. V. Guinea, and G. R. Plaza, 2019. Advances in micropipette aspiration: applications in cell biomechanics, models, and extended studies. Biophysical Journal 116:587–594.

67. Ko, K., S. R. Bandara, W. Zhou, L. Svenningsson, M. Porras-Gómez, N. Kambar, J. Dreher-Threlkeld, D. Topgaard, D. Hernández-Saavedra, S. Anakk, et al., 2024. Diet-induced obesity modulates close-packing of Triacylglycerols in lipid droplets of adipose tissue. Journal of the American Chemical Society 146:34796–34810.

68. Kumarage, T., S. Gupta, N. B. Morris, F. T. Doole, H. L. Scott, L.-R. Stingaciu, S. V. Pingali, J. Katsaras, G. Khelashvili, M. Doktorova, et al., 2025. Cholesterol modulates membrane elasticity via unified biophysical laws. Nature Communications 16:7024.

69. Almendro-Vedia, V. G., P. Natale, M. Mell, S. Bonneau, F. Monroy, F. Joubert, and I. López-Montero, 2017. Nonequilibrium fluctuations of lipid membranes by the rotating motor protein F1F0-ATP synthase. Proceedings of the National Academy of Sciences 114:11291–11296.

