## Supplementary material for "Ionizable Lipids Promote Curvature Remodeling and Altered Fluctuation Dynamics in Endosomal Membranes": SI

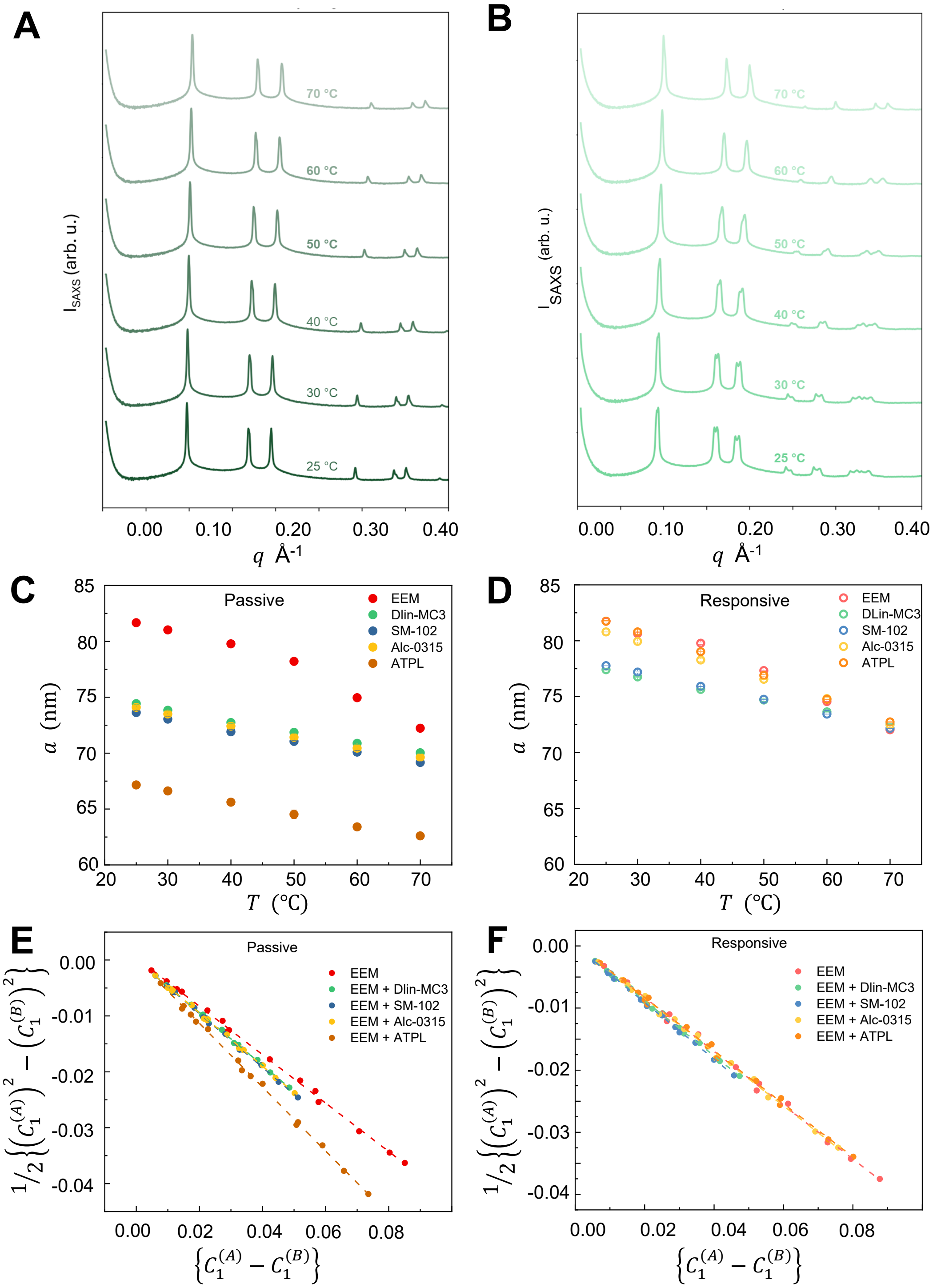


SI Figure 1: Spontaneous curvature calculations from SAXS for EEM with ILs/ATPL: (A) & (B) The SAXS profiles of the ($H_{\mathrm{II}}$) phase of DOPE doped with EEM + Dlin-MC3 at different temperatures, (C) & (D) The extracted lattice parameters of DOPE doped with EEM + IL/ATPL at different temperatures, (E) & (F) Linearized analysis of Eq 3 for DOPE doped with EEM +ILS/ATPL, for steady-state and responsive conditions respectively.

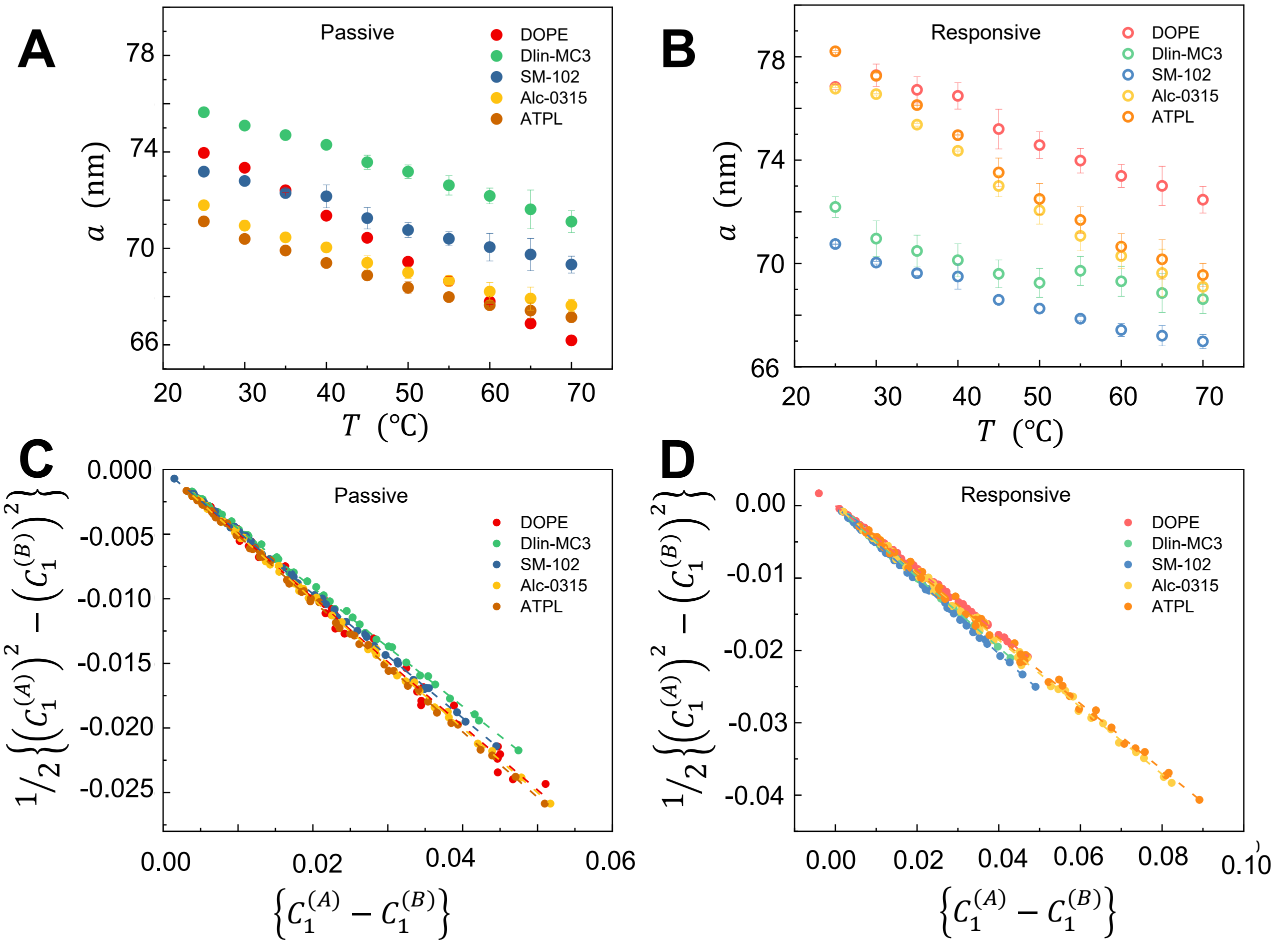


SI Figure 2: Spontaneous curvature calculations from SAXS for pure ILs/ATPL: (A) & (B) The extracted lattice parameters of DOPE doped with IL/ATPL at different temperatures, (E) & (F) Linearized analysis of Eq 3 for DOPE doped with ILS/ATPL, for steady-state and responsive conditions, respectively.

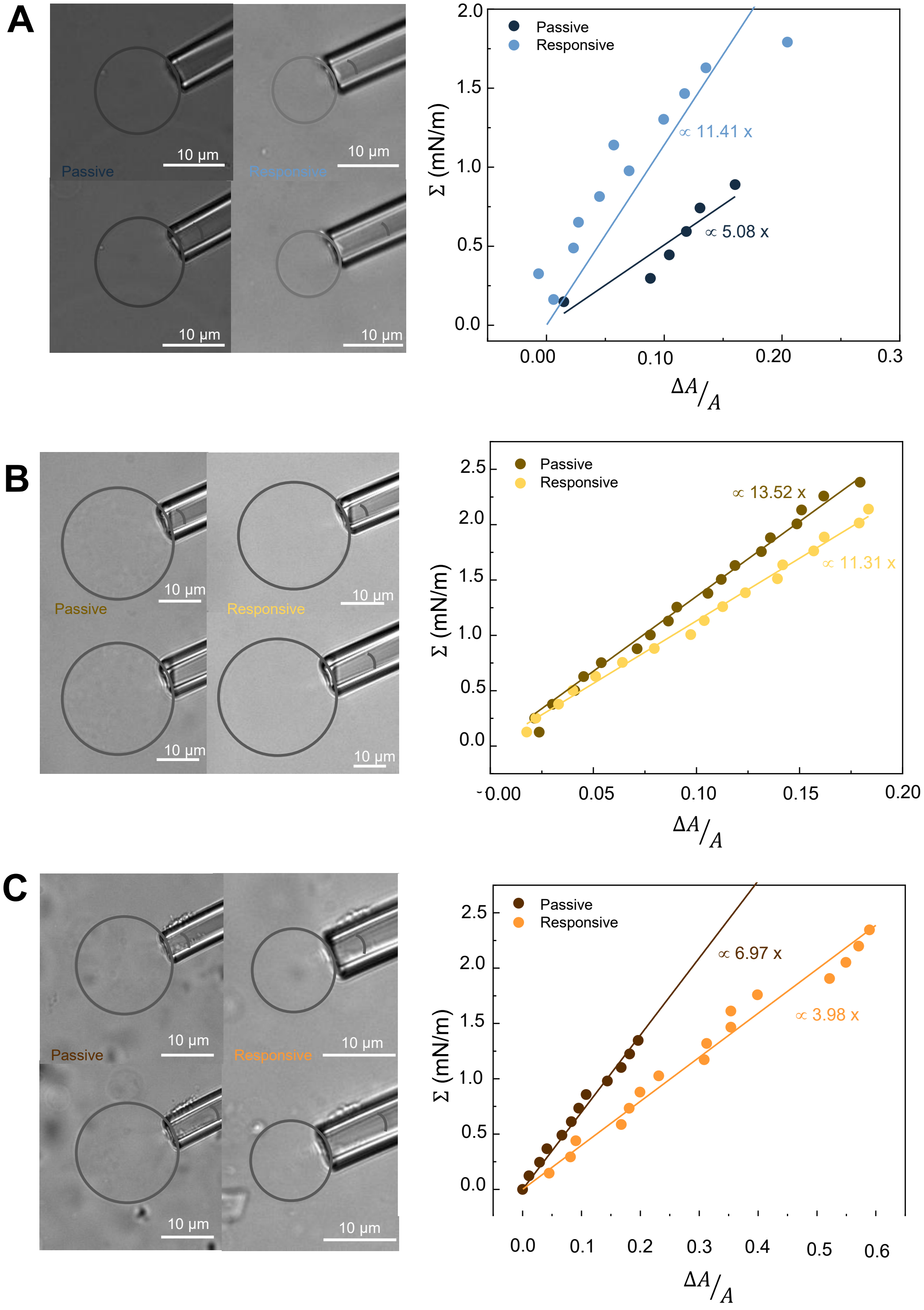


SI Figure 3: Micropipette aspiration analysis: Left panel shows 2D cross-sectional bright-field images of representative GUVs used for micropipette aspiration. The top image shows the vesicle prior to aspiration; the bottom image shows the change in aspirated membrane length induced by applied pressure, and the right panel shows the tension (Σ) vs. fractional change in surface area plot for the same GUVs for (A) SM-102, (B) Alc-0315 and (C) ATPL.

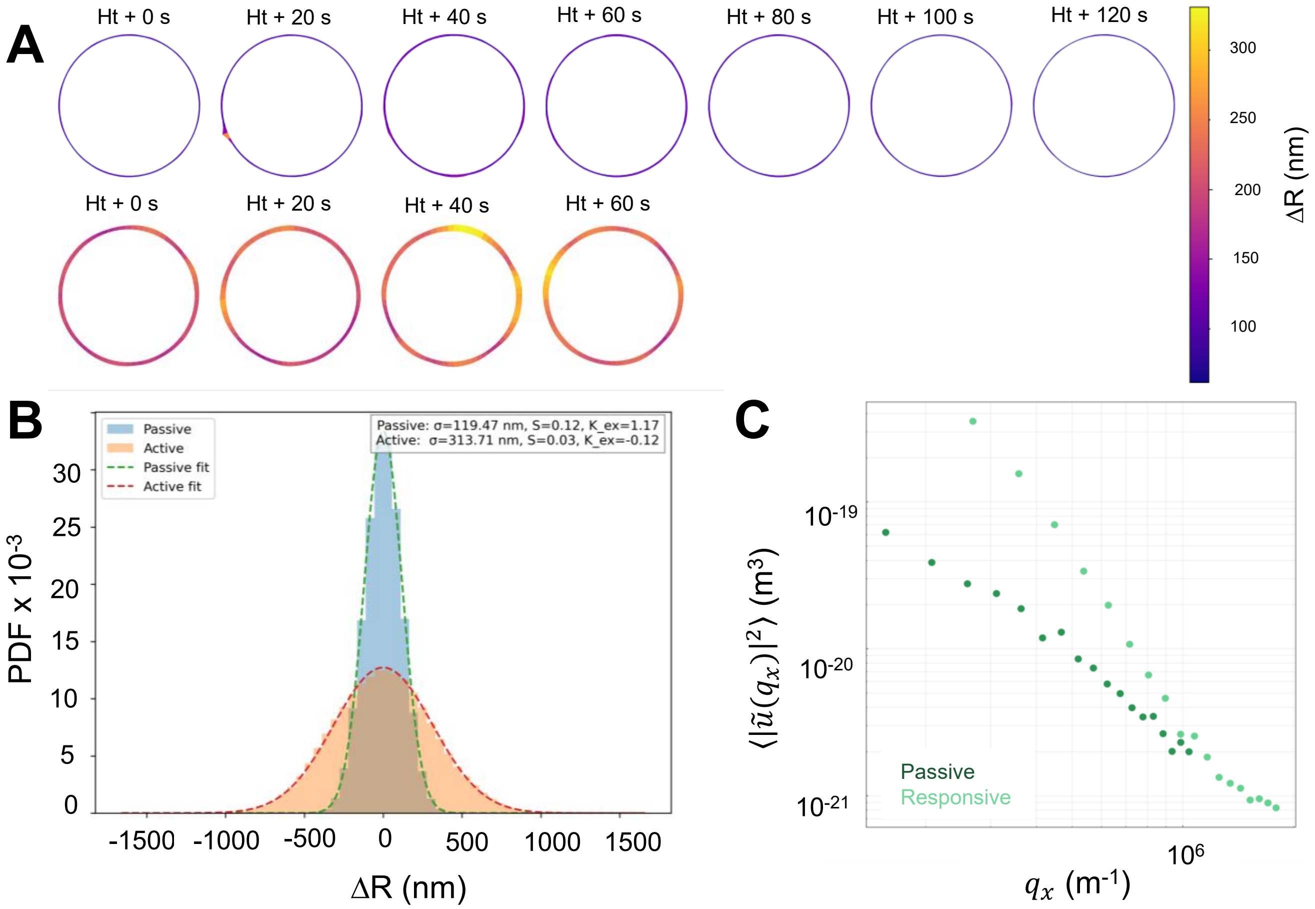


SI Figure 4: Vesicle fluctuation analysis of EEM+Dlin-MC3 GUVs: (A) Membrane fluctuation maps at different time intervals (top) steady-state and (bottom) responsive conditions. (B) Ensemble average PDF of GUVs and (C)experimental fluctuation spectra for the same vesicle under steady-state and responsive conditions.

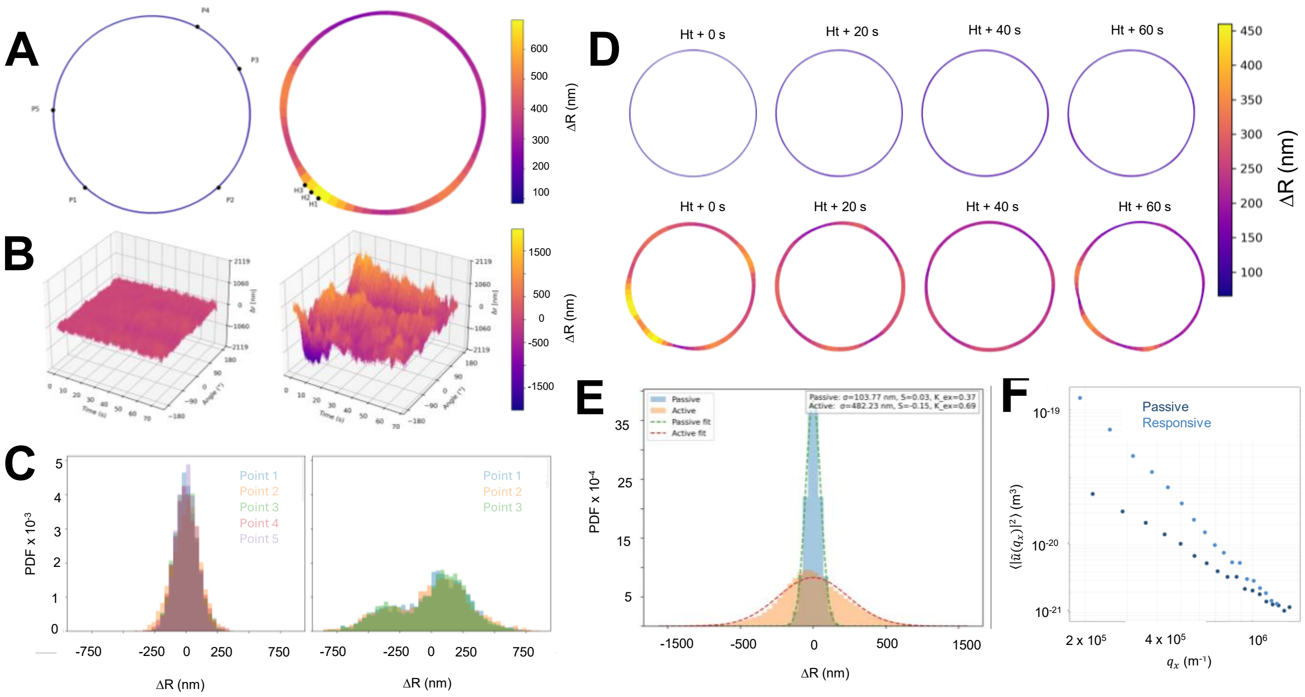


SI Figure 5: Vesicle fluctuation analysis of EEM+SM-102 GUVs: (A) Time-averaged fluctuation amplitude maps of vesicle contour, (B) Spatiotemporal maps of radial displacements, $\Delta R(\theta, t)$, as a function of angle and time, (C) Probability distribution functions (PDFs) of local radial displacements, $\Delta R$, sampled at multiple contour positions for steady-state conditions (left), and responsive membranes (right) conditions. (D) Membrane fluctuation maps at different time intervals (top) steady-state and (bottom) responsive conditions. (E) Ensemble average PDF of GUVs and (F)experimental fluctuation spectra for the same vesicle under steady-state and responsive conditions.

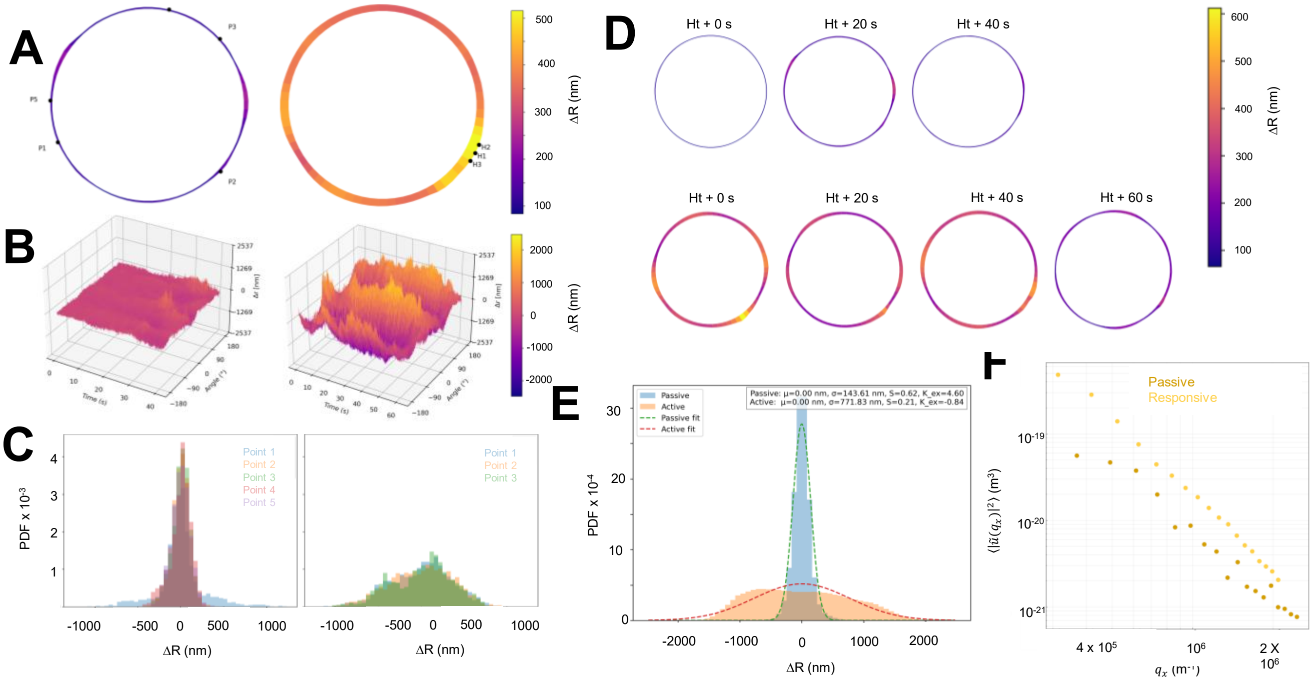


SI Figure 6: Vesicle fluctuation analysis of EEM+Alc-0315 GUVs: (A) Time-averaged fluctuation amplitude maps of vesicle contour, (B) Spatiotemporal maps of radial displacements, $\Delta R(\theta, t)$, as a function of angle and time, (C) Probability distribution functions (PDFs) of local radial displacements, $\Delta R$, sampled at multiple contour positions for steady-state conditions (left), and responsive membranes (right) conditions. (D) Membrane fluctuation maps at different time intervals (top) steady-state and (bottom) responsive conditions. (E) Ensemble average PDF of GUVs and (F)experimental fluctuation spectra for the same vesicle under steady-state and responsive conditions.

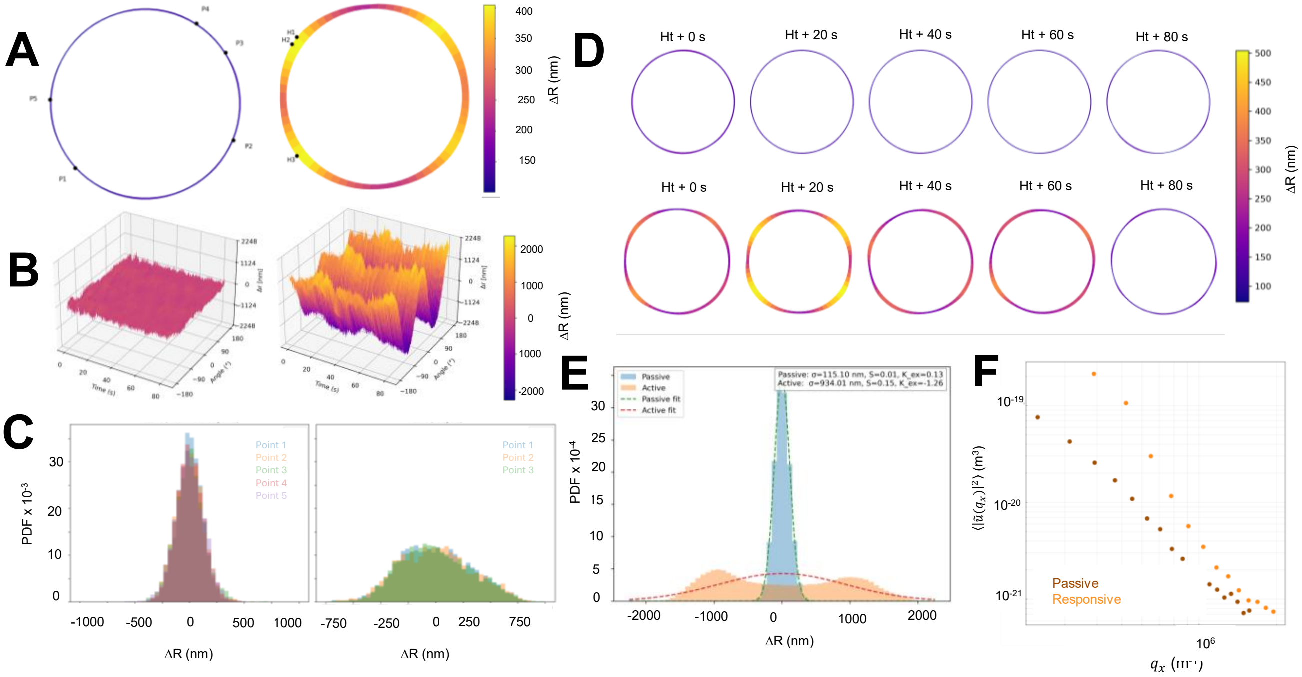


SI Figure 7: Vesicle fluctuation analysis of EEM+ATPL GUVs: (A) Time-averaged fluctuation amplitude maps of vesicle contour, (B) Spatiotemporal maps of radial displacements, $\Delta R(\theta, t)$, as a function of angle and time, (C) Probability distribution functions (PDFs) of local radial displacements, $\Delta R$, sampled at multiple contour positions for steady-state conditions (left), and responsive membranes (right) conditions. (D) Membrane fluctuation maps at different time intervals (top) steady-state and (bottom) responsive conditions. (E) Ensemble average PDF of GUVs and (F)experimental fluctuation spectra for the same vesicle under steady-state and responsive conditions.
